# Profiling genetically driven alternative splicing across the Indonesian Archipelago

**DOI:** 10.1101/2024.05.07.593052

**Authors:** Neke Ibeh, Pradiptajati Kusuma, Chelzie Crenna Darusallam, Safarina Malik, Herawati Sudoyo, Davis J. McCarthy, Irene Gallego Romero

## Abstract

One of the regulatory mechanisms influencing the functional capacity of genes is alternative splicing (AS). Previous studies exploring the splicing landscape of human tissues have shown that AS has contributed to human biology, especially in disease progression and the immune response. Nonetheless, this phenomenon remains poorly characterised across human populations, and it is unclear how genetic and environmental variation contribute to alternative splicing. Here, we examine a set of 115 Indonesian samples from three traditional island populations spanning the genetic ancestry cline that characterizes Island Southeast Asia. We conduct a global AS analysis between islands to ascertain the degree of functionally significant AS events and their consequences. Using a hierarchical event-based statistical model, we detected over 1,000 significant differential AS events across all comparisons. Additionally, we identify over 6,000 genetic variants associated with changes in splicing (splicing quantitative trait loci; sQTLs), some of which are driven by Papuan-like genetic ancestry, and only show partial overlap with other publicly available sQTL datasets derived from other populations. Computational predictions of RNA binding activity revealed that a fraction of these sQTLs directly modulate the binding propensity of proteins involved in the splicing regulation of immune genes. Overall, these results contribute towards elucidating the role of genetic variation in shaping gene regulation in one of the most diverse regions in the world.

## Introduction

Pre-mRNA splicing is a critical and highly regulated process through which multiple mRNA isoforms are produced from a single gene through the excision of introns and ligation of exons [1]. While constitutive splicing yields identically spliced mRNA isoforms, the process of alternative splicing (AS) produces isoforms that differ from each other based on their unique combinations of exons. AS is one of the regulatory mechanisms influencing the functional capacity of genes, and the resulting alternatively spliced isoforms contribute significantly to the protein diversity and functional complexity observed in eukaryotic organisms [2]. In humans, approximately 95% of multi-exon genes undergo AS [3, 4], and aberrant AS has been implicated in over 15% of human hereditary diseases and cancers [5, 6]. AS has been found to be highly specific, with isoform expression regularly restricted to certain tissues and cell-types [7–11], although the degree to which alternative isoforms are functional remains unclear [12–14]. Nevertheless, isoform-level transcriptome analyses have revealed that splicing can play significant roles in the cellular response to environmental cues, including immune pressures [15–21]. In humans, this natural variation in alternative splicing has been highlighted as a phenomenon influencing complex traits and disease prevalence [22].

Alternative splicing is tightly regulated through an intricate protein-RNA interaction network comprised of *cis* regulatory elements and *trans*-acting factors [23]. Genetic polymorphisms can alter these splicing regulatory elements and splice site usage, thereby influencing gene expression and protein products [24]. Splicing quantitative trait loci (sQTL) analysis has become a leading method for elucidating these genotype-splicing associations. sQTL analyses across multiple tissues have shed light on the contributions of AS to a number of traits, including breast cancer [25], Alzheimer’s disease [26], and schizophrenia [27]. The GTEx Consortium has recently characterised sQTLs in over 50 human healthy tissues [10], providing an overview of baseline healthy variation. Such studies are vital for advancing our understanding of gene regulation and disease mechanisms. They provide a bridge between genetic variation and complex traits, contributing to the development of innovative diagnostic tools and therapeutic interventions. The unprecedented scale and granularity of these analyses is, however, frustrated by the fact that the majority of participants involved in these studies are of either European or undocumented ancestry. This lack of diversity limits our collective understanding of variation in mRNA regulation and how these regulatory mechanisms might, in turn, contribute to human phenotypic diversity.

Multiple studies have sought to characterise the extent of variation across human populations in both splicing and the genetic mechanisms that regulate it, with existing data suggesting that both genetic ancestry and environmental variation make substantial contributions to these traits. In lymphoblastoid cell lines (LCLs) from 7 different global populations, population differences in genetic ancestry explained 25% of differences between individuals in genes expressing at least two transcripts [28], but expression differences, rather than splicing ones, accounted for the bulk of this effect. However, LCLs are known to trend towards a homogeneous pattern of gene expression that blunts the effect of inter-individual diversity [29], and thus this number may provide a lower-bound estimate of the true amount of variation. Indeed, the majority of AS events across GTEx tissues have been recently attributed to differences in ancestry between individuals, rather than to other demographic drivers such as age, sex or BMI [30].

The number of studies to map sQTLs is much more limited, and even more so when considering populations other than urban, genetically European cohorts; additionally most studies have used LCLs instead of primary tissue (reviewed in [31]). However, there are notable exceptions. A study of the response to infection in monocytes from individuals of European and African descent living in Belgium showed that genetic ancestry consistently contributed to differences in splicing, and highlighted the role of archaic introgression from Neanderthals [21] in particular. Recently, a separate study examining both expression and splicing in whole blood in East African populations has shown that overall, the genetic architecture of splicing is shared between individuals of African and European ancestry, but nonetheless identified a substantial number of sQTLs in individuals from Tanzania and Ethiopia that have not been reported in European cohorts [32].

Here, we focus on 115 previously described whole blood samples drawn from three traditional Indonesian populations [33, 34] that span the genetic ancestry cline that makes up the region [35]. We explore the prevalence and functional significance of alternative splicing variation across this region of the world, working towards deepening our understanding of the regulatory mechanisms that shape the landscape of immune gene regulation across the Indonesian archipelago. To elucidate the genetic variants influencing these splicing events, we identified 6,405 *cis*-sQTLs (4,199 unique SNPs) affecting 2,423 genes, and investigate possible biological mechanisms driving these observations. We discuss the implications of these gene regulatory patterns on a global scale, comparing our findings with those derived from European samples. Finally, we characterize the landscape of alternative splicing events across our groups of interest and investigate the functional consequences of the differential isoform usage between them. Thus, our work contributes to the growing catalog of putative regulatory elements that shape and influence alternative splicing in human populations.

## Materials and Methods

### RNA-seq processing and read alignment

We used a published set of 115 (all male) matched DNA and RNA data from three distinct Indonesian populations [33], available at (EGAS00001003671). The samples span the main west-east axis of genetic diversity in the Indonesian archipelago [35, 36]: (i) the inhabitants of Mentawai (MTW, n = 48), a small barrier island in western Indonesia, are of West Island Southeast Asian-like genetic ancestry; (ii) the Korowai (KOR, n = 19), from western New Guinea Island, are of Papuan-like genetic ancestry; while (iii) the inhabitants of Sumba (SMB, n = 48), a small island east of the Wallace line in central Indonesia, carry an approximately 80/20 admixture of either ancestry. Previous work has shown that the Korowai individuals in the data carry approximately 2% of archaic Denisovan introgression [34]. Both data collection and subsequent analyses were approved by the institutional review board at the Eijkman Institute (EIREC #90 and EIREC #126) and by the University of Melbourne’s Human Ethics Sub-Committee (approval 1851639.1). All individuals gave written informed consent for participation in the study. Permission to conduct research in Indonesia was granted by the Indonesian Institute of Sciences and by the Ministry for Research, Technology and Higher Education (RISTEK).

Reads were assessed with FastQC (v.0.11.9) [37] and pre-processed with Trimmomatic v0.36 [38], removing leading and trailing bases with a Phred score below 20 prior to any further analysis.

### Differential alternative splicing analysis

For event-based quantification of local splicing variation, we employed SUPPA2 [39]. SUPPA2 generates the percent spliced-in (PSI) values for each splicing event across all samples simultaneously using pseudoalignment-based transcript quantification. First, standard local splicing variations were computed using the SUPPA2 *generateEvents* command, applied to the reference annotation (GRCh38 Ensembl release 110). Then, the sample-wise PSI values for each event were calculated using transcript abundances (TPMs) obtained from Salmon v1.9.0 [40]. These PSI values denote the relative abundance of transcripts containing an exon (or intron, in RI cases) over the relative abundance of transcripts for the gene of interest containing the exon/intron. PSI scores were computed for five well-defined classes of alternative splicing events, namely, skipped exons (SE), retained introns (RI), mutually exclusive exons (MXE), alternative 5’ splice sites (A5SS), and alternative 3’ splice sites (A3SS). To ensure high confidence event calling, events were restricted to protein-coding and lincRNA genes, the percentage of exons with missing PSI values had to be below 5% per sample, and average TPM within a population had to be *>* 1).

To identify differential alternative splicing between the three groups, we corrected PSI values for batch effects and other technical confounders using fractional regression [41, 42]. Each splicing event was fit with logit-transformed PSI values (*glm* function in the R stats package (R v4.3.3), setting family = quasibinomial(‘logit’)). We selected the following covariates as the most relevant to incorporate into the model: RIN, sequencing batch, age of donor, and blood cell type proportions from previously computed [33] estimates of the proportion of CD8T, CD4T, NK, B cells, monocytes, and granulocytes in each sample. Using these corrected PSI values, differentially spliced events were identified by computing ΔPSI values between each pair of sample groups. Only splicing events that were detected in all groups were retained for differential testing (i.e., no NAs), as recommended in the SUPPA2 manual. Differential events were deemed statistically significant if they had a *|*ΔPSI*| ≥* 0.1 and FDR *<* 0.05.

In parallel, we conducted intron-based quantification of splicing variation (necessary for sQTL mapping) using LeafCutter [43], which identifies intron excision events across samples. Intron clustering (representing alternative intron excision events) was performed using default settings of 50 reads per cluster and a maximum intron length of 500kb. For each intron cluster, the proportion of reads supporting a specific intron excision event was calculated. Intron excision ratios were then standardized across individuals and quantile-normalized. LeafViz [43] was used for the annotation and visualization of intron clusters and splicing events.

### Comparative visualization of RNA-seq read alignments

We summarized exon- and junction-spanning RNA-seq densities for all sample groups using MISO’s [44] *sashimi plot* tool. Read densities for all exons were quantified using RPKM units [45]. Junction reads were plotted as arcs spanning the exons that the junction borders. Isoform structure was obtained from the GFF annotation (GRCh38 Ensembl release 110) of each splicing event. For ease of visualization, intron lengths were scaled down by factor of 30, and exon lengths were scaled down by a factor of 4.

### Isoform switching analyses

We identified instances of isoform switching by testing each individual isoform for differential usage across the three populations. Changes in isoform usage for a gene are represented by the difference in isoform fraction values (dIF), where IF = isoform expression/gene expression. Pseudoaligned transcript abundance and genomic coordinates were aggregated with the R package *IsoformSwitchAnalyzeR* [46]. Prior to isoform switch testing, we filtered out completely unused isoforms, single-isoform genes, as well as genes with an average TPM level (in each population) *<* 1. Confounding effects (RIN, sequencing batch, age, and blood cell type proportions, as above) were accounted for by applying the limma [47] *removeBatchEffect* function to the isoform abundance matrix. Statistical identification of isoform switches was conducted via *DEXseq* using default settings [48]. Isoforms were considered differentially switched and retained for further analysis if the difference in isoform fraction (dIF) *>* 0.1 and Benjamini-Hochberg-adjusted FDR *<* 0.05. We consider the first population in the switch comparison as the ground state, and the second population as the changed stated. For example, an up-regulated isoform in KOR versus MTW is one that is used more in MTW when compared to KOR. We then translated the coding sequences of the switching isoforms into amino acids, and predicted their coding capabilities, protein structure, peptide signaling, and presence of protein domain families using CPC2 [49], IUPred2A [50], SignalP [51], and Pfam [52], respectively.

The functional consequences of each significant isoform switch were evaluated by analyzing the annotation differences between the isoform(s) used more (switching up, dIF *>* 0) and the isoform(s) used less (switching down, dIF *<* 0). In other words, if a gene has at least one isoform with a significant change in usage between the three populations, the functional annotation of this isoform is compared to that of the isoform(s) with the compensatory change in usage. For genes with multiple significant switching events, we compared the functional annotation of all pairwise combinations of the isoforms involved. The following functional properties were compared: isoform coding potential, open reading frame (ORF) sequence similarity, the presence or absence of protein domain families, the presence or absence of signal peptides, the presence of intrinsically disordered regions (IDRs; regions that lack a fixed or ordered structure), and nonsense-mediated decay (NMD) sensitivity. For sequence similarity comparisons, we used the default minimum length difference cut-off of 10 amino acids [46]. Changes in protein domain or IDR length are only reported if the shorter protein domain (or IDR) is *<* 50% of the length of the longer region.

### *cis*-sQTL mapping

RNA-seq reads were aligned to GRCh38 Ensembl release 110 using the default settings of STAR (v.2.7) [53] with the exception of -alignEndsType EndToEnd to remove soft-clipping of the reads and -sdjbOverhang 100 for optimal splice junction overhang length. Mapped reads were used as input to LeafCutter [43] to obtain standardized and normalized intron excision ratios (the number of reads defining an excised intron over the total number of intron cluster reads), which were then used as phenotypes for sQTL mapping. We used QTLTools [54] to test for an association between variants and intron ratios within a *cis*-region of *±*1Mb of the intron cluster, using 10,000 adaptive permutations. After observing that the vast majority of our detected associations occurred within 250kb of the intron clusters, we restricted our analyses to this 250kb window. We controlled latent sources of variation using covariates identified with the Probabilistic Estimation of Expression Residuals (PEER) method [55]. The number of PEER factors was determined as a function of sample size, with the first 29 PEER factors (25% of our sample size) selected. The top 5 genotyping principal components were included to account for population structure, since they account for approximately 9% of the genotype variance observed, with diminishing returns for all subsequent PCs. Nominal *p*-values for each variant-phenotype pair were obtained by testing the alternative hypothesis that the slope of the linear regression model between genotype and excision ratios deviates from 0.

### Identifying shared *cis*-sQTLs and differences in effect sizes

We compared our Indonesian sQTLs to two European whole blood sQTL studies: GTEx v8 [10] (n = 670) and BLUEPRINT [9] (n = 197) with freely available complete summary statistics. For both European studies, the sQTL mapping approaches that were employed are in line with those used for our Indonesian data here, making comparisons possible without extensive raw data reprocessing. Specifically, both studies applied a linear modeling approach, testing for associations with variants within *±*1Mb of each gene’s transcription start site. For consistency, GTEX and BLUEPRINT sQTL sets were restricted to 250kb, and coordinates for the BLUEPRINT data were converted from hg19 to hg38 using the R package *liftOver* v1.26.0 [56]. Shared and population-specific sQTLs across Indonesian and European populations were assessed using the multivariate adaptive shrinkage (mash) approach implemented in the R package *mashr* [57]. The input for mash consisted of sQTL effect sizes and their standard errors (obtained from the QTLTools output). The correlation structure among the null tests was estimated using a large, random subset of all tests (40%). For each intron cluster within each gene, the SNP with the smallest *p*-value across all tested SNPs was retained in order to produce a confident set of sQTLs. The data-driven covariance matrix was constructed using this strong set, and posterior mean effect sizes were calculated by applying the mash model that was built using the random set. sQTLs were considered shared across populations if the magnitude of effect sizes was within a factor of 0.5 between groups, and the sign of the effect was the same. A local false sign rate (LFSR) *<* 0.05 was used as a threshold for significance.

### Variance in sQTL genotype explained by local genetic ancestry

Using all significant permutation-based sQTLs, we quantified the variance in sQTL genotype explained by modern local genetic ancestry (LA). To do this, we adapted a previously described approach [58], fitting a linear model V = *α ×* PAP + *β* for each variant. V is the genotype vector and PAP is a covariate which represents the number of alleles assigned to Papuan-like genetic ancestry. This analysis was conducted using the 73 samples (30 Mentawai, 29 Sumba, 14 Korowai) with available 30x depth whole-genome sequencing data [34]. Variants were categorized as highly correlated with LA if they had an absolute *R*^2^ *>* 0.7.

### Gene ontology overrepresentation analysis

We tested for overrepresentation of GO and KEGG terms using the R package *clusterProfiler* (v.4.0.5) [59], setting the gene universe as all tested genes. We used an FDR threshold of 0.05 to identify significantly enriched terms. For GO term representations across sGenes, we employed REVIGO [60] (parameters: allowed similarity = 0.9, database =*H.sapiens*, semantic metric = SimRel) to remove highly redundant GO terms from *clusterProfiler* output and visualize semantic similarity-based GO term representations.

### Estimation of sQTL variant pathogenicity

We intersected our sQTL SNPs with clinically annotated variants in ClinVar (https://www.ncbi.nlm.nih.gov/clinvar) in order to assess their possible pathogenicity. We considered two sets of variants for analysis—lead SNPs located within the gene body, and all sQTL SNPs that were located directly within a splice junction.

### Evaluating the impact of sequence variations on the binding affinity of splicing RNA binding proteins

We used DeepCLIP to quantify the effects of SNPs on protein-RNA binding [61]. Specifically, we used DeepCLIPs pre-trained models for 33 splicing-associated RNA binding proteins (RBPs). These RBPs were chosen because they have been previously characterized in eCLIP (enhanced Cross-Linking and ImmunoPrecipitation) studies from HepG2 and K562 cell lines and have well-documented roles in splicing regulation and spliceosome activity [62, 63]. Using the pre-trained models, we ran prediction in paired sequence mode using all significant sQTL SNPs, with 10 bp flanking sequences on both sides. The output from DeepCLIP is a set of binding profiles and overall scores for the reference and variant sequences. The binding profile scores range from 0 to 1, and indicate whether regions of a sequence contain potential binding sites (1) or if they are most likely to be random genetic background (0) [61].

### Colocalization analysis with GWAS hematological traits

Colocalization analyses were performed between sQTLs and 13 hematological traits using coloc v5.2.3 [64]. We tested for colocalization between our detected permutation-based sQTLS and GWAS loci from a trans-ethnic study that included East Asian populations [65]. We assumed a prior probability that a SNP is associated with i) the GWAS trait (p1, default = 1 *×* 10*^−^*^4^), ii) alternative splicing (p2, default = 1 *×* 10*^−^*^4^), and iii) both the GWAS trait and alternative splicing (p12, default = 1 *×* 10*^−^*^6^). We identified robust colocalization with the default threshold of CCV *>* 0.8 and a ratio CCV/DCV *>* 5. The 13 hematological traits measured in the East Asian populations were: basophil count, eosinophil count, hematocrit, hemoglobin concentration, lymphocyte count, mean corpuscular hemoglobin, mean corpuscular hemoglobin concentration, mean corpuscular volume, monocyte count, myeloid white blood cell count, neutrophil count, platelet count, and RBC density. All GWAS summary statistics were obtained from the GWAS catalog. Since we did not have access to linkage disequilibrium values for the Indonesian samples and our sample size is too small to build a robust LD matrix [66], the colocalization analyses performed here did not allow for the identification of multiple causal variants.

## Results

### Characterizing differential alternative splicing events between Indonesian populations

To characterize the splicing landscape across the Indonesian archipelago, we carried out alternative splicing analyses using paired-end RNA-seq data generated from 115 previously described male whole blood samples [33]. Samples were obtained from three Indonesian populations: the people of Mentawai (of genetically West Island Southeast Asian-like ancestry), the people of Sumba (approximately 80/20% admixed between West Island Southeast Asian-like and Papuan-like genetic ancestries), and the Korowai of New Guinea Island (of Papuan-like genetic ancestry) (Figure 1A).

**Figure 1.**
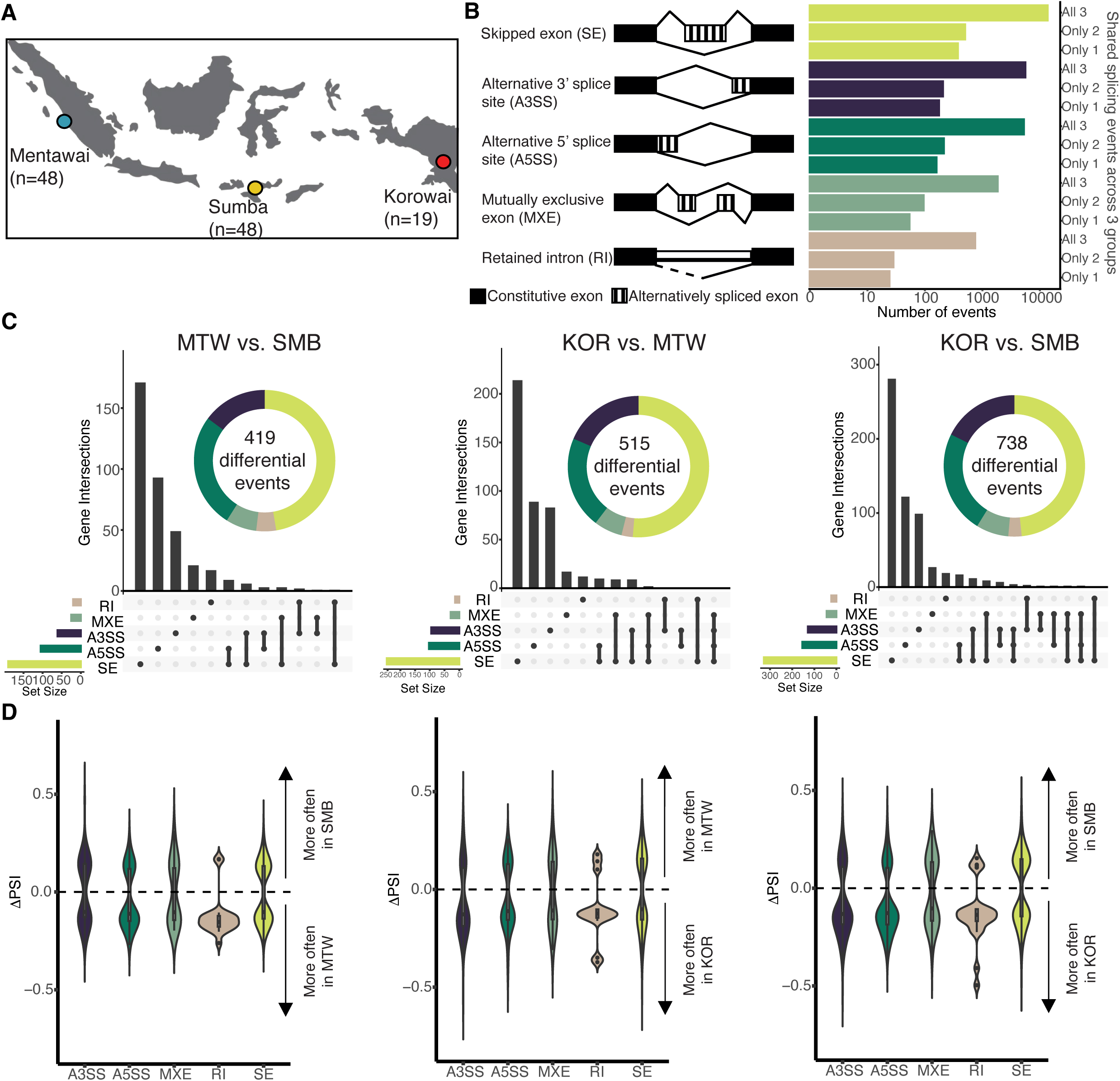
Characterization of differential alternative splicing across Indonesian populations. **A**) Geographical location of the three sampling sites (n = 115). **B**) Schematic diagram of the five alternative splicing events profiled with SUPPA2. Horizontal bar plots correspond to the event categories to the left, and illustrate the sharing of alternative splicing events across the three island groups (i.e., the splicing event either occurs in all three groups, in only two of the three groups, or in only one group). The x-axis is *log*_10_ transformed. **C**) UpSet plots summarize the DAS gene intersections for the three pairwise comparisons. Only significant events (*|*ΔPSI*| ≥* 0.1, FDR *<* 0.05) are shown. **D**) For each population pair, the distributions of ΔPSI values are plotted for each splicing category (A3SS, A5SS, MXE, RI, and SE). The direction of the arrows indicate whether the alternatively spliced exon is excluded more or included more in each sample (i.e. exon exclusion levels). In the case of an MXE event, if the strand is positive, then the inclusion form includes the first exon and excludes the second; if the strand is negative, then the inclusion form includes the second exon and skips the first.

Across the three populations, we observed 38,611 alternative splicing events in the Korowai, while the Mentawai and Sumba groups had 39,728 and 39,769 alternative splicing events, respectively. This slight difference in total detected events can be attributed to the smaller sample size of the Korowai group (n = 19). Our results show that the splicing events across these three populations were generally shared. Specifically, 38,144 (94.2%) of all alternative splicing events were seen in all three island groups, 1,337 events (3.3%) were shared between only two of three groups, and 1,002 events (2.5%) were found in just one of the three groups (Figure 1B). We additionally observe that the Mentawai and Sumba groups make up the majority of the pairwise AS event sharing (74.1% of all events shared by any two island groups, Supplementary Figure 1B-F), although this is likely attributable to our increased power in these two groups relative to the Korowai. To gain further insight into the genes that contain these 1,002 population-specific alternative splicing patterns (434 events in MTW, 447 events in SMB, and 121 events in KOR), we tested for their enrichment against GO and KEGG pathways (Table S2). Overlapping enriched GO categories and KEGG pathways for the Korowai population are related to nervous system function, transmembrane transportation, and substance dependence. In Mentawai, significantly enriched GO categories are generally related to regulation of synaptic signaling, however, no enriched KEGG pathways were detected. For Sumba, enriched GO categories and KEGG pathways include renal system processes, hormone transport, and calcium signaling.

To better understand the splicing variation across these populations, we conducted a differential alternative splicing analysis. We identified 419 significant (*|*ΔPSI*| ≥* 0.1 and FDR *<* 0.05) differential alternative splicing events between Mentawai and Sumba, 515 events between Korowai and Mentawai, and 738 events between Korowai and Sumba (Figure 1C and D). These differences are in line with previously reported gene expression patterns across these groups, with the Korowai vs. Sumba comparison yielding the most differentially expressed genes, closely followed by Korowai vs. Mentawai [33]. These findings suggest that the Korowai population might be driving a lot of the observed inter-island variability, with the Sumba and Mentawai populations exhibiting higher levels of shared homogeneity, although we cannot confidently disambiguate whether these differences are caused by genetic or environmental factors. Differentially alternatively spliced (DAS) genes between Korowai and Sumba are related to mRNA processing (e.g., *SLTM*, *BARD1*, *DYRK1A*, *CDK11A*, *QKI*, *RNPS1*), splicing regulation (e.g., *RBM23*, *TRA2B*, *TMBIM6*, *MBNL1*, *HNRNPH1*, *RNPS1*,

*PTBP2*), and cell cycle signaling (e.g., *MRE11*, *BRIP1*, *DONSON*, *ZWILCH*, *CDC14B*), indicating potential population-based differences in post-transcriptional regulatory patterns (Table S3). While DAS genes between Korowai and Mentawai were not enriched for either GO or KEGG terms, we found that DAS genes between Mentawai and Sumba were enriched for transcription corepressor (e.g., *DPF2*, *ATF7IP*, *MTA1*, *HDAC9*, *SF1*) and coactivator (e.g., *MED24*, *KMT2C*, *SMARCA2*, *MED12*, *ACTN1*) activity (Table S3). Overall, these findings indicate that the DAS genes between these populations are predominantly impacting aspects of cell division and regulation of gene expression.

### Characterisation of Indonesian *cis*-sQTLs in whole blood

To understand the genetic regulation of alternative splicing across our Indonesian populations, we performed a *cis*-sQTL mapping analysis using all 115 samples. sQTLs were mapped using linear regression to quantify the association between the intron excision ratios and SNP genotypes (Materials and Methods). Splice ratios from a total of 49,122 splice clusters (i.e., overlapping introns that share a spice donor or acceptor site) were included in the sQTL analysis. We detected a total of 6,077 *cis*-sQTLs affecting 1,977 genes at an FDR level of 0.01, comprising 3,658 unique SNPs (Table S4 and Table S5). Positional enrichment of significant sQTL SNPs showed clustering closest to the detected splice junctions, with 75.2% of SNPs occurring within 50kb of the nearest junction (Figure 2A). This observation is consistent with previous research suggesting that most variants that fall in close proximity to splice junctions influence splicing regulatory functions [67, 68] GO enrichment analysis of sGenes (genes with a significant sQTL) revealed enrichment for numerous immune-related pathways, including: immune response-regulating signaling (e.g., *THEMIS2*, *ERMAP*, *GBP2*, *VAV3*, *FCRL3*, *AIM2*, *FCGR3A*), immune response-activating signaling (e.g., *DENND1B*, *MAPKAPK2*, *CR1*, *NLRC4*, *NAGK*, *IFIH1*, *PRKCD*), phagocytosis (e.g., *VAV3*, *NCF2*, *C4BPA*, *DYSF*, *MARCO*, *CD302*), regulation of innate immune response (e.g., *GBP2*, *ADAR*, *FCRL3*, *IFI16*, *AIM2*, *CFH*), and interleukin-1 beta production (e.g., *IFI16*, *AIM2*, *NLRC4*, *CASP8*, *GHRL*, *CX3CR1*), highlighting the impact of splicing on immune-related genes (Supplementary Figure 3, Table S6). Indeed, some of the strongest sQTL signals are present in immune-related genes such as *TRIM58*, which is additionally implicated in malignancies such as lung cancer, liver cancer, and pancreatic ductal adenocarcinoma [69–72], and is involved in innate immunity and cell proliferation [69, 73], and *ERMAP*, which is a B7 family immune regulator that has been found to promote the phagocytosis of tumor cells [74–76] and encodes the protein responsible for the Scianna blood group system [77] (Supplementary Figure 4).

**Figure 2.**
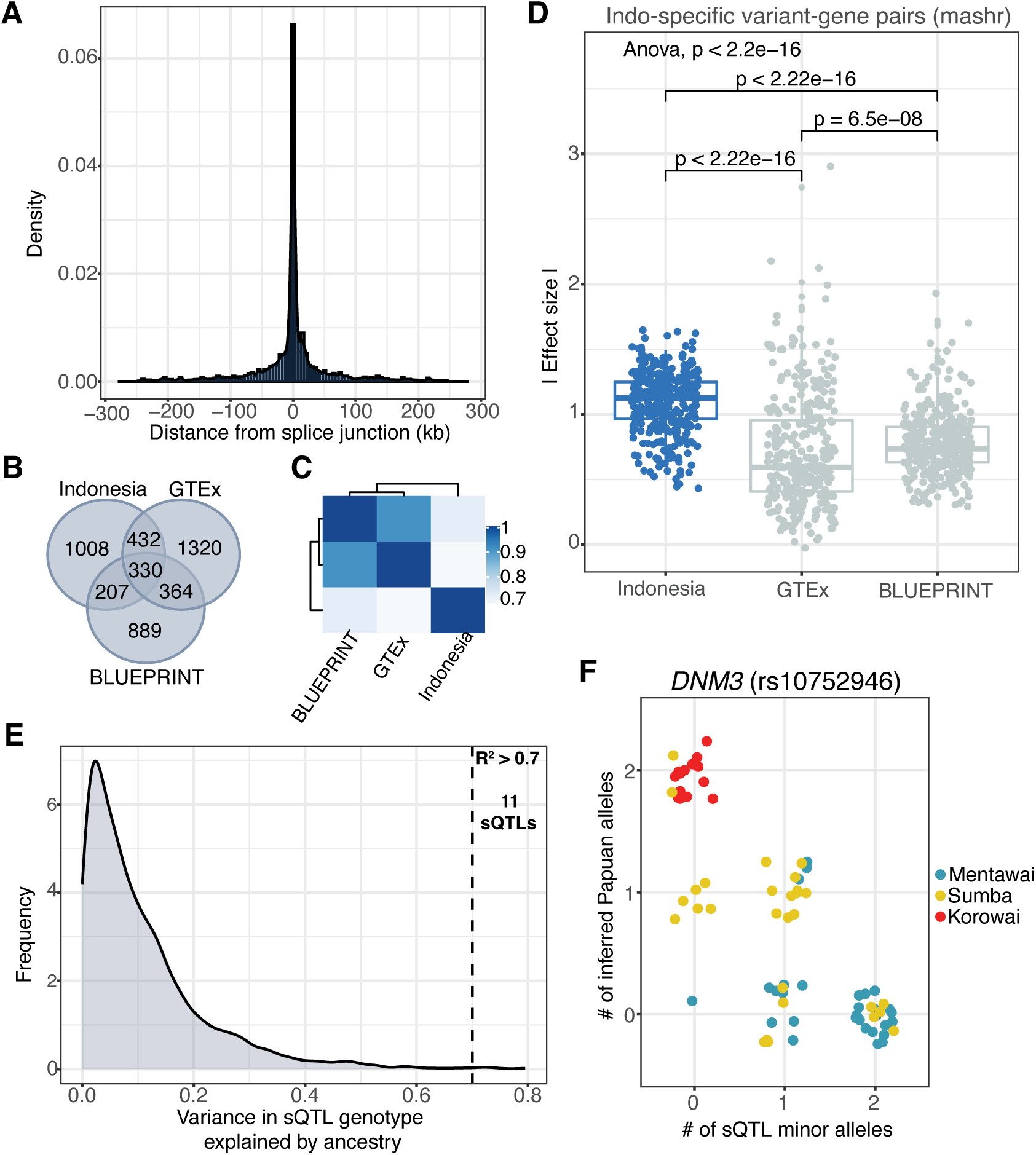
Indonesian *cis*-sQTL characteristics. **A**) Relative distance of sQTLs to the nearest splice junction. **B**) Shared and unique sGenes across the Indonesian and European datasets. **C**) Pairwise sharing of sQTLs between the datasets evaluated by mashR. **D**) Distribution of effect sizes for Indonesia-specific sQTLs. **E**) Linear regression between the number of QTL minor alleles and number of inferred Papuan alleles reveals 11 sQTLs largely driven by Papuan-like genetic ancestry (*R*^2^ *>* 0.7). **F**) An sQTL that is strongly correlated with Papuan-like genetic ancestry. Plot shows the relationship between the number of inferred Papuan alleles and sQTL minor alleles.

Given that the majority of sQTL studies to date have profiled European populations, we wanted to compare the overlap between our detected sQTLs and previously published datasets. We focused our comparison on two large studies of whole blood, namely: GTEx v8 [10] (n = 670) and BLUEPRINT [9] (n = 197). We found that sQTL effects are largely shared between the two European data sets, but they are not shared to the same extent with the Indonesian data. Out of 4,550 unique sGenes (amongst the three datasets), 330 (7.3%) are shared across all datasets, 364 are detected in both European cohorts but not in the Indonesian data (8%), and 1008 (22.2%) are solely in the Indonesian data; similar values were private to both GTEx and BLUEPRINT, suggesting either incomplete power to detect sQTLs across studies, winner’s curse, or a genuine poor generalizabilty of sQTLs across studies (Figure 2B). To better estimate shared and population-specific effects and ascertain the degree of sharing between our detected Indonesian sQTLs and the European sQTLs, we applied a multivariate adaptive shrinkage model, mashr. Using a local false sign rate cut-off of 0.05, an sQTL is considered shared if the effect sizes share the same sign and fall within a factor of 0.5 of each other. As with the more näıve approach above, sQTL effects are largely shared across European datasets (Figure 2C). Additionally, we detected 312 Indonesia-specific (LFSR *<* 0.05) SNP-sGene pairs (Figure 2D). In line with previous eQTL analyses carried out on this set of samples [34], we observe significantly larger effect sizes of Indonesian-specific sQTLs within Indonesia than we do in the European datasets (ANOVA, *p <* 2.2e-16; Figure 2D), again suggestive of incomplete power across all considered datasets, and highlighting the importance of establishing larger cohorts for QTL mapping in general.

While there was no evidence of enrichment of GO or KEGG terms across these Indonesia-specific genes, many genes with sQTLs are implicated in signal regulation (i.e. *IRF1*, *SIRPB1*) and oncogene pathways (i.e. *RAP1A*, *RAP1B*, *FES*, and *DNM3*). In spite of our limited sample size, these findings suggest a distinctive and distinguishable (relative to previously characterized European data) Indonesian sQTL signal, as we would expect sharing between GTEx and BLUEPRINT to also be negatively impacted by the relatively low sample size in BLUEPRINT and their distinct processing pipelines. By profiling the differences and similarities across these groups, we can more accurately characterize the gene regulatory mechanisms underlying diversity in genetic traits, highlighting the value of trans-ethnic, trans-environment QTL mapping to maximize discovery of gene regulatory variation across humans [30, 32].

After quantifying sQTL-level differences between the Indonesian and European cohorts, we wanted to assess the extent to which variance in sQTL genotype is driven by local genetic ancestry in modern Indonesians. Specifically, using local genetic ancestry haplotype information for each significant sQTL, we investigated whether there was a correlation between the inferred ancestral source of the genotype and gene splicing (Materials and Methods). We found 11 sQTLs that show strong evidence (*R*^2^ *>* 0.7) of being driven by Papuan-like genetic ancestry (Figure 2E, Table S7). Although small and likely underpowered due to the very limited representation of Papuan-like genetic ancestry in our dataset, this set included several immune genes, one of which was *DNM3* (Figure 2F). Otherwise known as Dynamin-3, *DNM3* is a microtubule-associated gene that plays a fundamental role in membrane trafficking and the regulation of vesicle formation in cells [78], and has been implicated in various functions related to the development and colony formation of megakaryocytes [79–81]. Human *DNM3* contains 21 exons, with at least 5 of its exons being alternatively spliced to generate several isoforms [82]. The Papuan-driven sQTL event that we identified for *DNM3* is a skipping event of exon 17 (*β* = 0.55, *q*-value = 3.5 *×* 10*^−^*^4^), associated with the SNP rs10752946, which lies within intron 20 of *DNM3* (Figure 2F, Figure 3A). The canonical isoform for this gene, ENST00000627582.3, includes exon 17 and encodes a protein made up of 863 amino acids (GRCh38 Ensembl release 110). Conversely, isoform ENST00000367731.5 is the product of the alternative splicing event that excludes exon 17, resulting in a loss of 4 amino acids which do not appear to have a significant impact on protein structure predictions made with AlphaFold. Our results indicate that the Korowai samples have a higher rate of exon 17 exclusion/skipping (22.5%) than the Mentawai (7.6%) and Sumba (7.5%) groups; in other words, a higher proportion of expressed *DNM3* isoforms in the Korowai group exclude exon 17.

**Figure 3.**
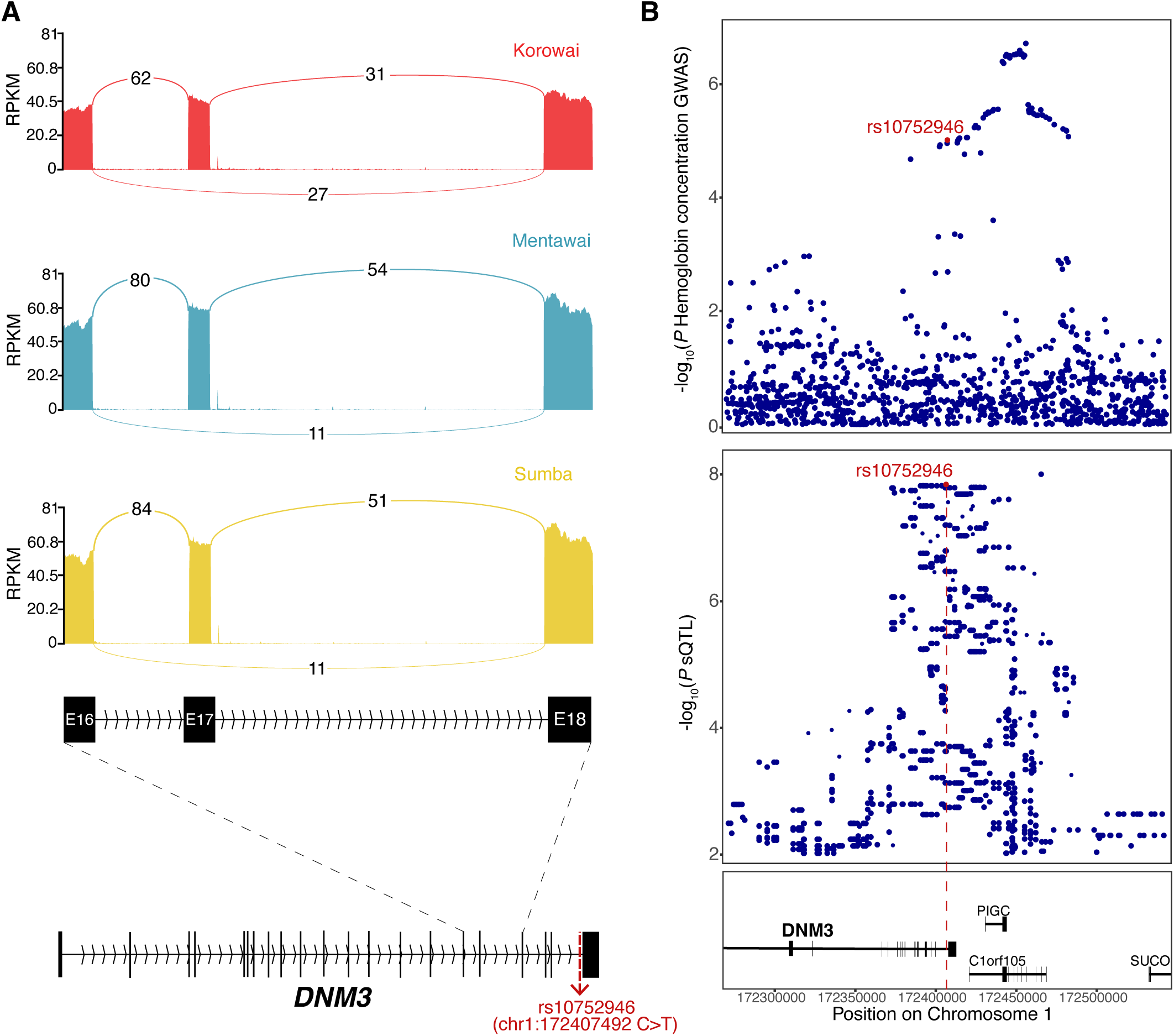
Splicing regulation in the *DNM3* gene. **A)** Sashimi plot illustrating the skipped exon event (exon 17) in the *DNM3* gene. Isoform expression was summarized using the MISO framework [44]. The numbers over each curved line represent the junction spanning reads. The canonical isoform for *DNM3* is provided. The labels E16, E17, and E18 represent exons 16, 17, and 18, respectively. **B**) Statistically significant GWAS colocalization between the LAI-driven sQTL in the *DNM3* gene (rs10752946) and the GWAS hemoglobin concentration trait [65] measured in East Asian populations.

### Indonesian whole blood *cis*-sQTLs colocalize with hematological GWAS traits

To shed light on the potential interplay between gene splicing regulation and complex traits, we sought to explore whether the Indonesian sQTLs colocalized with any hematological GWAS traits. Using coloc, a Bayesian test for genetic colocalization, we performed colocalization analyses between all significant sQTLs and 13 blood-related traits using summary statistics from a trans-ethnic study which included 151,807 East Asian participants [65]. Overall, we identified 45 unique sGenes (3.4 % of all significant sGenes) that colocalized with 12 different traits, from a total of 68 significant GWAS trait-sGene pairs (File S1). Seven of these genes colocalized with three traits simultaneously. Of these, *FCGRT*, which encodes the neonatal Fc receptor (FcRn) colocalized with red blood cell density, hemoglobin concentration, and hematocrit, while *MAEA*, a macrophage erythroblast attacher, colocalised with mean corpuscular hemoglobin, mean corpuscular volume, and red blood cell density. We also found that the local genetic ancestry-driven sQTL at *DNM3* significantly colocalizes with hemoglobin concentration (H4 = 0.837, H4/H3 = 8.445), a trait which refers to the amount of hemoglobin protein present in a specific volume of blood (Figure 3B).

### Assessing the pathogenicity of *cis*-sQTL variants

Given the growing awareness of the role of splicing in genetic disease and the ancestral diversity biases in existing sQTL datasets, we intersected sQTL SNPs with clinically annotated variants on ClinVar. We considered two sets of SNPs: i) all lead sQTL SNPs within the gene body and ii) all sQTL SNPs falling within splice junctions. Across both SNP sets, the analysis yielded 120 total hits, with 116 of these having benign/likely benign status, 3 variants of uncertain significance (VUS), and 1 variant with a GWAS disease trait classification (Table S8). The 3 VUS occur in *KLRC4* (rs139613925, in-sample alternative allele frequency 0.481), *HADHB* (rs2303893, in-sample alternative allele frequency 0.366), and *DENND3* (rs2289001, in-sample alternative allele frequency 0.307). The protein encoded by *KLRC4* belongs to the Natural Killer Group 2 (NKG2) family of receptors. These receptors are primarily expressed on natural killer (NK) cells and some subsets of T cells, where they play important roles in the recognition and regulation of immune responses [83]. The *HADHB* gene product, MTP-beta, plays a central role in the breakdown of long-chain fatty acids within mitochondria [84]. *DENND3* plays a role in regulating intracellular vesicle trafficking, and has been implicated in cancer, where dysregulation of vesicle trafficking pathways can contribute to tumor progression and metastasis [85,86]. The GWAS variant, rs2763979 (in-sample alternative allele frequency 0.468), occurs in *HSPA1B* and has reported associations with cancer (*p*-value = 2 *×* 10*^−^*^14^, European population, n = 475,312) [87] and systemic lupus erythematosus (*p*-value = 6 *×* 10*^−^*^6^, Thai population, n = 4,088) [88].

### *cis*-sQTL SNPs influence splicing-RBP binding affinity

We used a deep-learning convolutional neural network, DeepCLIP, to predict the effect of our sQTLs on RBP-RNA binding as a possible mechanism of action. We employed DeepCLIPs pre-trained models for 33 splicing-associated RNA binding proteins (RBPs) with well-documented roles in splicing regulation and spliceosome activity [62], and assessed the impact on their binding abilities across all 3,658 significant lead SNPs within 250kb of the nearest splice site, adding 10bp flanking sequences on both sides. For each reference-alternative allele pair and each RBP, we obtained binding profiles indicating the likelihood of RBP binding (for the respective protein) along the RNA sequence. These binding affinity scores, ranging from 0 to 1, can be interpreted as the strength of RBP binding [61]. Globally, we observe that most sQTL SNPs do not significantly increase or decrease RNA binding affinity, and their difference scores (reference score - alternative score) tend to fall within the range (99% CI) of likely values (Figure 4A, Supplementary Figure 5). However, for RBPs with very uniform motifs, such as EWSR1 (Ewing sarcoma protein) [89], the introduction of sQTL SNPs has a marked effect on the sequence binding scores, yielding large differences in predicted binding affinity between the reference and the alternative allele (Supplementary Figure 5).

**Figure 4.**
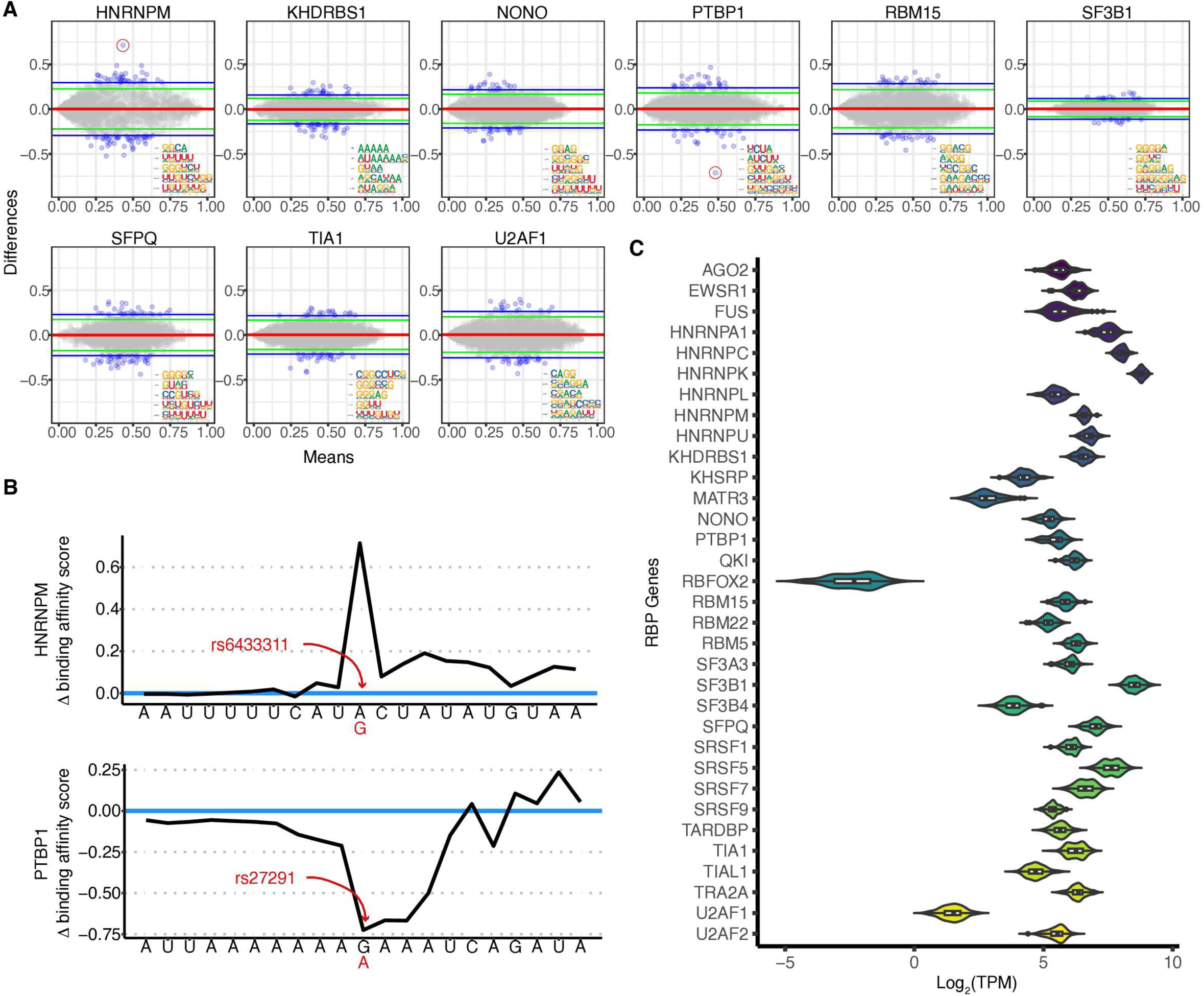
sQTL impact on RBP binding affinity. **A)** Mean-difference plots for 9 of the 33 splicing-RBPs. Each point represents a pair of 21bp-long sequences centered on the lead sQTL SNP (ref/alt alleles). The y-axis depicts the difference between the DeepCLIP reference sequence score and the DeepCLIP variant sequence score, while the x-axis depicts the average between these two scores. Horizontal green and blue lines represent the 95% and 99% CIs, respectively. Points that are outside of the 99% CI are colored blue. CNN filters predicted by DeepCLIP, ranked by score, are provided for each protein. **B)** DeepCLIP binding profiles for HNRNPM and PTBP1. The variant-reference sequence pairs in question are circled in panel A. **C)** Normalized expression (TPM) of the 33 chosen splicing-associated RNA binding proteins.

Conversely, we also observe RBPs with only one or two outliers, suggesting a biologically significant impact of the sQTL SNP on RBP activity (Figure 4A and B). Specifically, for the HNRNPM protein, which is moderately expressed within our dataset (Figure 4C), we predict a 0.71 increase in binding affinity at rs6433311 (A*>*G, Figure 4B), which is an sQTL for the *DYNC1I2* gene; in our data, the G allele at rs6433311 is associated with an effect size of −1.174 relative to the A allele (indicating exon exclusion, Supplementary Figure 6A), and segregating at a frequency of 0.739. The predicted increase in HNRNPM binding as a consequence of this A*>*G intronic variant is consistent with previous findings that HNRNPM recognizes GU-rich cis-elements, primarily in introns [90–92]. HNRNPM is a member of the heterogeneous nuclear ribonucleoprotein (hnRNP) family involved in RNA processing, including pre-mRNA splicing and mRNA transport [93], and has been found to promote exon skipping across various experimental conditions [92, 94–96]. The *DYNC1I2* gene, as a component of the dynactin complex [97], may indirectly influence RNA transport within the cell, potentially interacting with HNRNPM-associated mRNA complexes during transport. In addition, we predict a 0.73 decrease in the binding affinity of *PTBP1* at rs27291 (G*>*A, Figure 4B), which is an sQTL for the *LNPEP* gene; in this case, the A allele at rs27291 in our data is associated with an effect size of 0.672 (indicating exon inclusion) and segregating at an allele frequency of 0.434. PTBP1, also known as Polypyrimidine Tract Binding Protein 1, plays crucial roles in alternative splicing, mRNA stability, mRNA localization, and translation initiation [98–101]. Containing four highly conserved RNA binding domains that recognize short pyrimidine-rich sequences [102], this splicing factor regulates alternative splicing by inducing exon skipping [103–105]. Thus, a decrease in the binding activity of this protein may result in higher exon inclusion, which is in line with the positive effect size that we observe at this splicing junction (Supplementary Figure 6B).

### Genome-wide isoform switching analysis reveals significant changes in functional protein domains as a consequence of fluctuations in isoform usage

In order to better assess the functional implications of our detected splicing events, we conducted an isoform switching analysis across each of the three pairwise group comparisons (Mentawai vs. Sumba, Korowai vs. Sumba, and Korowai vs. Mentawai). Isoform switching is defined as the change in relative abundance of different isoforms from the same gene between conditions. Isoforms were considered differentially switched and retained for further analysis if the difference in isoform fraction (ΔIF) *>* 0.1 and FDR *<* 0.05. In total, we identified 165 significant isoform switches across 110 genes between Korowai and Mentawai, 244 switches across 158 genes between Korowai and Sumba, and 100 switches across 66 genes between Mentawai and Sumba(Table S9 and Table S10). Altogether, these events impacted 239 unique genes and 389 unique isoforms. Then, to ascertain the potential biological impact of these isoform switches, we tested for changes in the functional properties of the transcripts at the sequence level. These properties included the coding potential of the transcripts, the functional domains and signal peptides (that are either present or absent), the presence of intrinsically disordered regions, or IDRs (regions that lack a fixed or ordered structure), the sensitivity of the transcript to nonsense-mediated decay (NMD), and the similarity of the open reading frame (ORF) sequences between transcripts. For every detected isoform switch in a given gene, the annotations for the most-used isoform (ΔIF *>* 0.1) and the least-used isoform (ΔIF *<* 0.1) were compared. Differences in the annotations between the up-regulated isoform and the down-regulated isoform were categorized as isoform switches with predicted functional consequences (Figure 5A).

**Figure 5.**
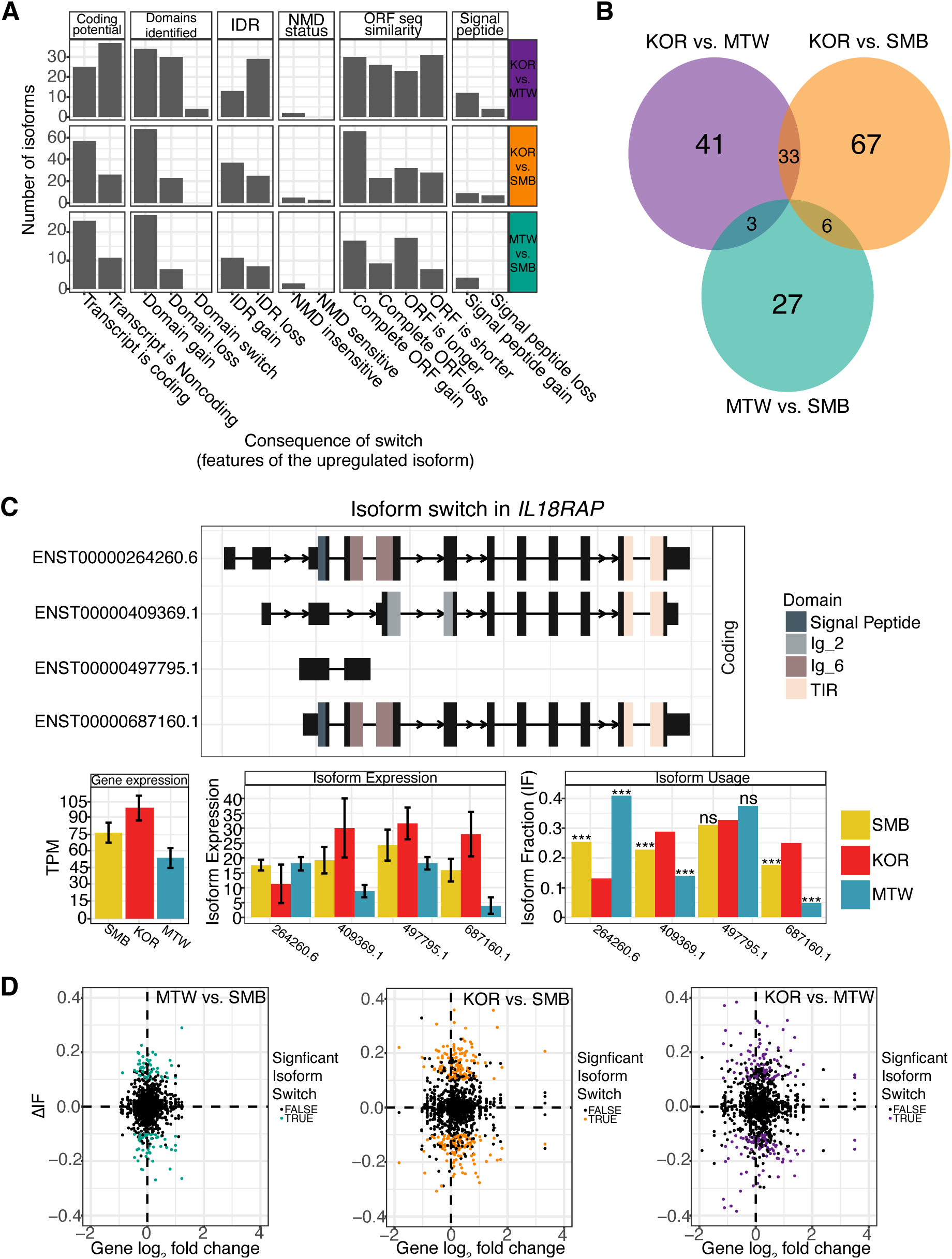
Global isoform switching analysis and functional consequences. **A**) For each pairwise group comparison, the consequences of all significant isoform switching events are described. Specifically, the annotation differences between the up-regulated isoform (dIF *>* 0) and the down-regulated isoform (dIF *<* 0) are summarized. The first population in the switch comparison is the ground state, and the second population is the changed stated. Thus, an up-regulated isoform in the KOR vs. MTW comparison, for example, is one that is used more in MTW relative to KOR. **B**) Venn diagram of the number of genes with overlapping isoform switching events and functional consequences (FDR *<* 0.05) across all pairwise comparisons. **C**) The isoform switch in *IL18RAP*. Exons with annotated domains are coloured, and those without are black boxes. Normalized *IL18RAP* expression (TPM), isoform expression, and isoform usage are plotted. Error-bars indicate 95% confidence intervals. Level of significance (with KOR as the reference) is denoted by the asterisks. **D**) Changes in isoform switches against changes in gene expression (x-axis) for each pairwise comparison. Significant (ΔIF *>* 0.1 and FDR *<* 0.05) isoform switches are coloured accordingly.

Across all switching events between the three Indonesian populations, the most frequent functional consequences of isoform switching were changes in ORF sequence similarity (36.7%), protein domains identified (23%), and coding potential (21.4%, File S2-File S5). When focusing on switches between Korowai and either Mentawai or Sumba, isoforms expressed at higher levels in the Mentawai or Sumba groups were more likely to contain an additional domain, with 14.8% of all switching events resulting in a protein domain gain in the up-regulated isoform, while 7.1% resulted in a domain loss in the up-regulated isoform (Figure 5A). Furthermore, for up-regulated isoforms in the Sumba population (KOR vs. SMB and MTW vs. SMB), a greater number of switches resulted in the increased usage of a coding transcript (15%) as opposed to a non-coding transcript (6.8%). Interestingly, when compared to the Korowai population, isoforms up-regulated in the Sumba population are more likely to have an IDR gain (9.2%) rather than an IDR loss (5%) and are also more likely to be coding, whereas the inverse is true for isoforms unregulated in the Mentawai population versus the Korowai population (Figure 5A). IDRs act as a flexible point of contact for protein-protein interactions and are often enriched in protein products that play a critical role in signaling, regulation, and molecular recognition [106–108].

We also found that the number of these functionally significant switches, and the extent of overlap between pairwise comparisons, varied greatly. For the Mentawai vs. Sumba comparison, we detected statistically significant isoform switching events with a functional consequence across 36 genes, 106 genes for Korowai vs. Sumba, and 77 genes for the Korowai vs. Mentawai comparison (58.3%, 68.8%, and 73.2% of all isoform switches in each comparison; Figure 5B). The overlap in functionally significant switching genes was greatest between the pairs involving the Korowai group, with the Mentawai vs. Sumba comparison sharing very few switching genes with either of the other two pairs. Moreover, there were no functionally significant switching genes that were shared across the three sets.

Many of the significant isoform switching events occurred in genes that play critical roles in the immune response, one of which is the *IL18RAP* gene, the most significant isoform switching event across all comparisons (*q*-value = 5 *×* 10*^−^*^27^, Table S9). The IL18RAP protein (Interleukin 18 Receptor Accessory Protein) serves as a receptor accessory protein for interleukin-18 (IL-18), a cytokine involved in immune responses and inflammation [109]. As such, *IL18RAP* is a crucial component of the IL-18 receptor complex, and the dysregulation of *IL18RAP* expression has been implicated in various diseases, including autoimmune disorders, inflammatory conditions, and cancer [110]. In our data, we observe preferential usage of the longer *IL18RAP* isoform (ENST00000264260.6) in the MTW group (isoform switch *q*-value = 1.8 *×* 10*^−^*^8^), while the SMB and KOR groups exhibit preferential usage of the shorter isoforms ENST00000409369.1 and ENST00000687160.1 (Figure 5C). Notably, the ENST00000264260.6 isoform contains a signal peptide at the third exon, while isoforms ENST00000409369.1 and ENST00000497795.1 do not. Signal peptides play a key role in directing proteins to their appropriate cellular locations. Thus, higher usage of the *IL18RAP* signal peptide-containing isoforms (ENST00000264260.6 and ENST00000687160.1) might be indicative of an up-regulation of IL18RAP protein translocation. Furthermore, isoforms ENST00000264260.6, ENST00000409369.1, and ENST00000687160.1 contain immunoglobulin domainsIg 2 (PF13895), and Ig 6 (PF18452), which play essential roles in antigen recognition, cell adhesion, receptor-ligand interactions, and structural stability [111]. Higher isoform usage of these isoforms might therefore influence structural stability and protein diversity.

In addition, we looked for evidence of significant isoform switching across our 11 detected Papuan ancestry-driven sGenes, such as *DNM3*. We observed statistically significant isoform switching that also resulted in functional domain changes in the *BST1* gene (Supplementary Figure 7A). *BST1* (bone marrow stromal cell antigen 1) is an immune gene that facilitates pre-B-cell growth and regulates leukocyte diapedesis in inflammation [112]. The isoform switch in the *BST1* gene reveals increased usage of the ENST00000265016.9 isoform within the SMB group (isoform switch *q*-value = 1.3 *×* 10*^−^*^4^), while KOR samples exhibit a higher isoform fraction of the ENST00000514989.1 isoform. The ENST00000514989.1 isoform is non-coding, which means that a switch in isoform usage results in a loss of coding potential. Interestingly, the ENST00000265016.9 isoform contains a rib hydrolase domain at exons 1-8, which is essential for *BST1* enzymatic activity as a NAD+ glycohydrolase [113].

### Most differentially expressed genes are not differentially spliced

Our isoform switching analyses illustrate how critical alternative splicing is for the diversification of gene expression at both the mRNA and protein levels (Figures 5A and D). To assess the extent to which splicing and transcriptional regulation are linked across each pairwise comparison, we intersected the differentially spliced genes with previously reported [33] differentially expressed genes in the same groups. We found that differentially spliced genes were very rarely differentially expressed, with only 13 (Fishers exact test *p* = 9.7e-04), 65 (Fishers exact test *p* = 1.1e-07), and 39 (Fishers exact test *p* = 2.1e-06) shared genes across both tests for the Mentawai vs. Sumba, Korowai vs. Sumba, and Korowai vs. Mentawai comparisons, respectively (Supplementary Figure 7B). In other words, for most genes, we observe changes in the mRNA splice variants produced without a reciprocal change in total expression. These results support previous findings of independent regulation of gene splicing and expression [22, 114–117].

## Discussion

It is well-established that alternative splicing influences various biological functions [2,7–11,22]. Studies of the link between genotypic and splicing diversity in humans, however, have not yet sampled the breadth of genetic and environmental diversity that characterizes our species, making it challenging to assess the degree to which population-specific forces contribute to splicing variation. In this work, using a set of 115 samples from three traditional island populations spanning the regions genetic ancestry cline, we characterize the global alternative splicing landscape across these populations and identify genetic variants that are associated with splicing (sQTLs), thereby providing a comprehensive map of genetically regulated alternative splicing events in human whole blood.

Our detection of alternative splicing revealed that the most frequent splicing event types were skipped exon events (50.1%) and the least frequent were retained intron events (3.5%). While these proportions are expected in higher eukaryotes [41, 118–123], it is important to note that short-read algorithms for detecting AS events are less likely to misassemble skipped exon events and can therefore identify them with high accuracy, but they suffer from poor precision with respect to RI detection [124, 125]. Accurately disambiguating real splicing events from background noise (such as partial transcript processing) is greatly facilitated by long-read sequencing technologies, which enable much more precise discovery and reconstruction of full-length isoforms, significantly improving the characterization of functional transcripts [126]. Thus, for future isoform-specific alternative splicing studies, the application of long-read technologies should be prioritized.

We conducted a global differential alternative splicing analysis between these groups and detected over 1500 significant events across all comparisons, with Mentawai and Sumba exhibiting the lowest number of differential events (i.e., the highest level of alternative splicing similarity). Given that these two populations share very similar proportions of West Island Southeast Asian-like genetic ancestry, while the Korowai are of Papuan-like genetic ancestry, our results suggest that ancestral differences might play a role in driving some of the variation in splicing across these populations. Interestingly, our *cis*-sQTL mapping analysis revealed 11 sQTLs driven by local genetic ancestry differences between groups. Furthermore, we detected population-specific patterns of splicing regulation at a global scale, identifying 312 Indonesia-specific SNP-sGene pairs with no evidence of shared effects with European sQTLs. Although our analysis is limited by our sample size, these findings suggest that alternative splicing variation between populations reflects a complex interplay between genetic and environmental factors.

As a means of characterizing the phenotypic consequence of alternative splicing variation across these Indonesian samples, we investigated whether GWAS signals for hematological traits measured in East Asian participants colocalized with Indonesian sQTLs. Of the 13 GWAS traits examined, 45 unique sGenes showed strong evidence of colocalisation with 12 distinct traits. Although colocalization analysis alone is not a sufficient means by which causal genes can be defined, the Papuan-driven splicing gene *DNM3* significantly colocalized with hemoglobin concentration, indicating its potential role in regulating this trait, and is notable observation given the role of high altitude corridors in the settlement of New Guinea Island [127].

To further bridge the gap between our identified sQTLs and their functional consequences, we employed a deep-learning model, DeepCLIP, to predict changes in RBP binding affinity. We found that the DeepCLIP predictions of the impact of sQTL SNPs on RBP binding dynamics correlated with the direction of effects that were observed at these splice junctions. In particular, we predicted a potential increase in HNRNPM binding affinity at rs6433311 (A*>*G, *DYNC1I2*), while our sQTL mapping analysis showed that the G allele at rs6433311 is associated with an effect size of −1.174 relative to the A allele. Leafcutter-predicted skipping junctions (exon skipping events) yield negative effect sizes, while inclusion junctions (exon inclusion events) yield positive effect sizes. Therefore, the predicted increase in binding affinity of a splicing factor that promotes exon exclusion [92, 94–96] should be associated with a negative effect size at this SNP, which is indeed what we observe. For another splicing factor, PTBP1, DeepCLIP analysis highlighted a potential binding disruption at the rs27291 G*>*A mutation in the *LNPEP* gene. In our data, the A allele of this sQTL SNP is associated with an effect size of 0.672, which is indicative of increased exon inclusion. Given the role PTBP1 plays in suppressing exon inclusion (thereby promoting exon exclusion) [103–105], the predicted reduction in PTBP1 activity directly correlates with a positive effect size for this sQTL and a higher exon inclusion rate for individuals harbouring the rs27291 A allele. While these results are consistent with previous findings that sQTLs disrupt the binding affinity of core and auxiliary splice proteins [22, 128], it is important to note that we have not conducted any experimental validation of the predicted protein binding activity, thereby limiting our ability to evaluate the accuracy of the deep-learning predictions. Nonetheless, our results highlight the molecular mechanisms through which genetic variants impact splicing, and demonstrate the value of applying deep learning methods to refine our collective understanding of the consequences of human genetic variation.

We also explored evidence of functionally significant isoform switching events across the three populations, identifying frequent changes in protein domains (domain gain/loss), NMD status, and ORF sequence similarity. We observed the greatest number of isoform switching genes in comparisons involving the Korowai population, in line with the population-level alternative splicing differences we identified with SUPPA2. Notably, we report that when an isoform switch occurs between the Korowai group and the Sumba or Mentawai groups, the up-regulated isoforms in Sumba or Mentawai are more likely to: i) contain an additional protein domain, ii) contain a signal peptide, and iii) gain a complete open reading frame. Our analysis identified several significant isoform switching events in genes critical for the immune response, including the *IL18RAP* and *BST1* genes. In the *IL18RAP* gene, we observed preferential usage of different isoforms across populations, with potential implications for protein translocation and structural stability. Similarly, in the *BST1* gene, isoform switching resulted in changes in coding potential and the presence of functional domains, highlighting the importance of alternative splicing in immune regulation and disease processes.

As previous studies have reported [129–131], we find that a large subset of genes exhibit isoform switches without significant changes in their global expression levels. Such genes would typically be overlooked in RNA-seq studies that strictly focus on differential gene expression, thereby limiting critical biological insight. Irrespective of the global expression levels of a gene, changes in the relative expression of its isoforms influence protein abundance, which in turn modulates cellular processes. Furthermore, we detect minimal overlap between differentially expressed genes and differentially alternatively spliced genes, indicating that the genetic control of splicing and transcription are independent, as previously observed [17, 21, 22, 30, 132–134]. Indeed, across our Indonesian dataset, splicing and expression affect different pathways, as DE and DAS genes were enriched for distinct biological processes and molecular functions. DAS genes were mostly involved in processes related to post-transcriptional regulation, DNA replication, and cell-cycle signaling. In contrast, DE genes were enriched in regulatory pathways related to the adaptive immune response and nervous system function [33]. Together, this study provides a comprehensive catalog of genetically regulated alternative splicing events in whole blood Indonesian samples, and adds to our knowledge of genomic and regional drivers of gene regulatory variation across the Indonesian archipelago.

## Acknowledgements

We thank all of the original study participants and the members of the Genome Diversity and Disease team, without whom this work would not have been possible, along with members of the Gallego Romero and McCarthy labs for comments. P.K is supported by a Wellcome International Training Fellowship (no. 222992/Z/21/Z). St Vincent’s Institute acknowledges the infrastructure support it receives from the National Health and Medical Research Council Independent Research Institutes Infrastructure Support Program and from the Victorian Government through its Operational Infrastructure Support Program.

## Data and code availability

All data used in this project was previously published and is deposited at the European Genome Archive under accession EGAS00001003671. Genome-wide significant and nominal test statistics for all sQTL tests are provided as supplementary tables and are publicly available on Figshare: https://melbourne.figshare.com/projects/Profiling_genetically_driven_alternative_splicing_across_the_Indonesian_Archipelago/204111. Analysis code is available at https://gitlab.svi.edu.au/igr-lab/indo_splicing

## Supplementary Figures

**Supplementary Figure 1.**
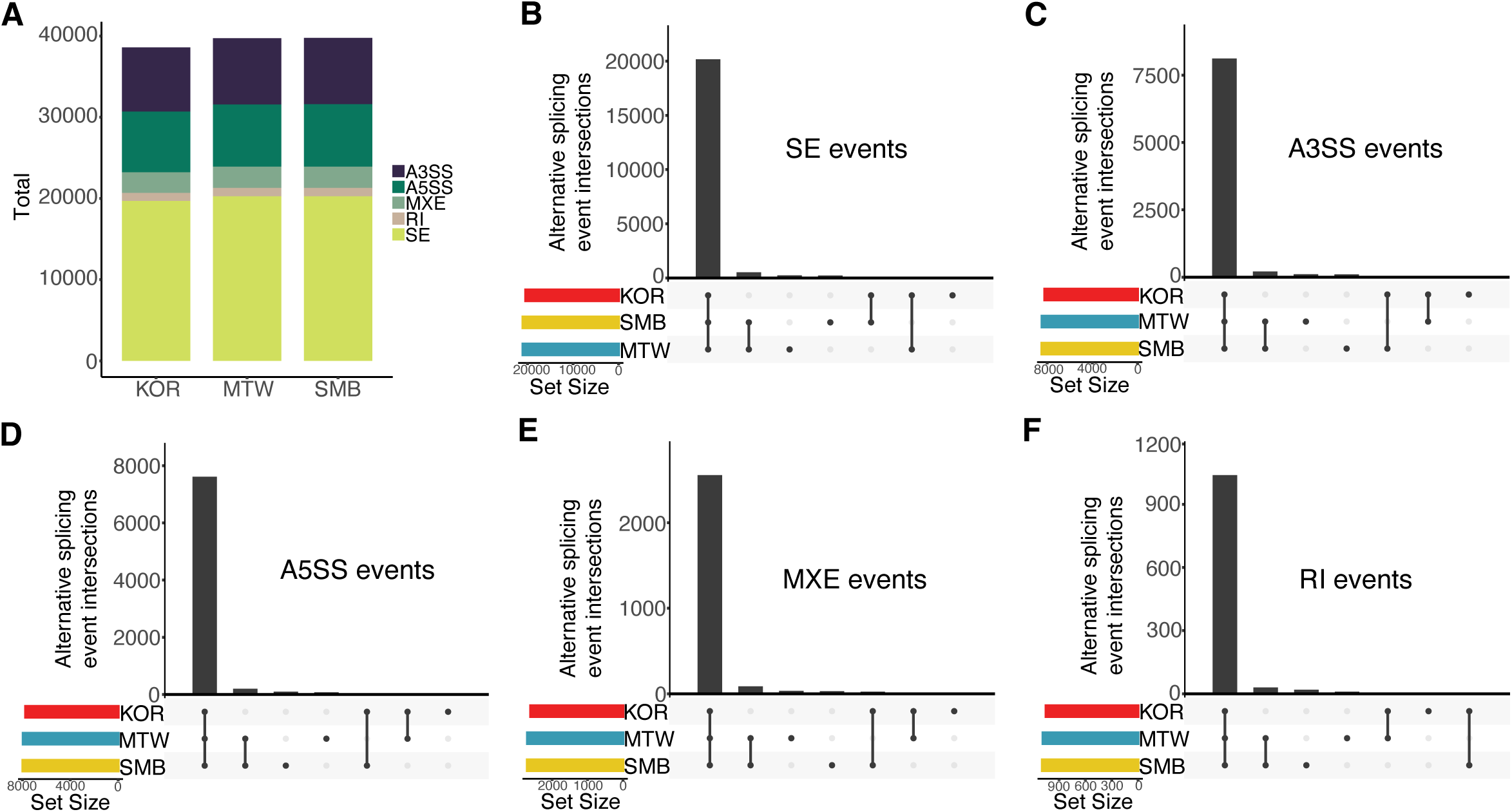
AS events quantified with SUPPA2. **A)** Proportion of each AS event type for Korowai, Mentawai, and Sumba. **B)-F)** AS event intersections between sample groups.

**Supplementary Figure 2.**
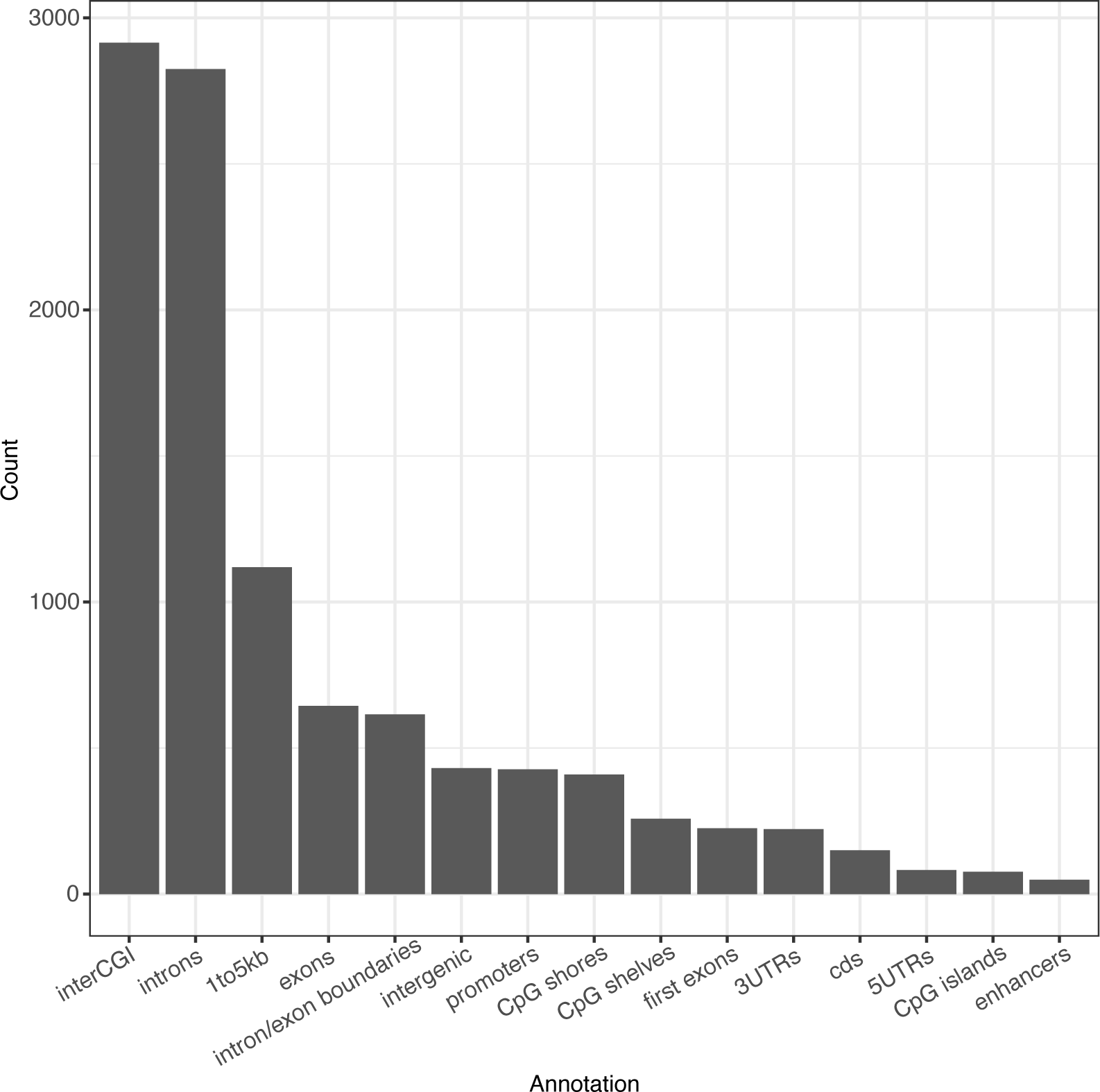
Genomic locations of sQTLs. Significant sQTLs and their prevalance in annotated regions.

**Supplementary Figure 3.**
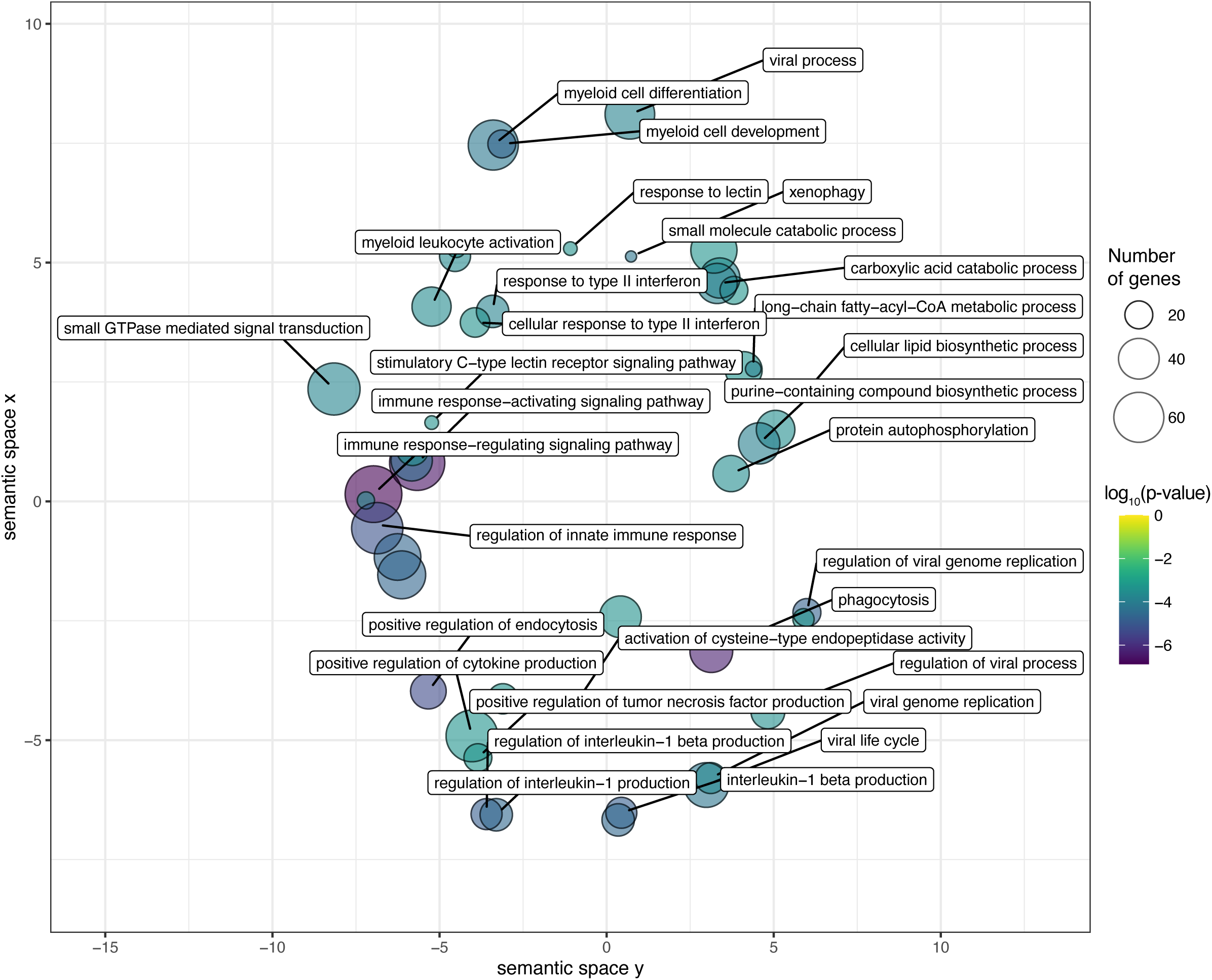
GO term clustering of differentially alternatively spliced genes using semantic similarity measures in Revigo.

**Supplementary Figure 4.**
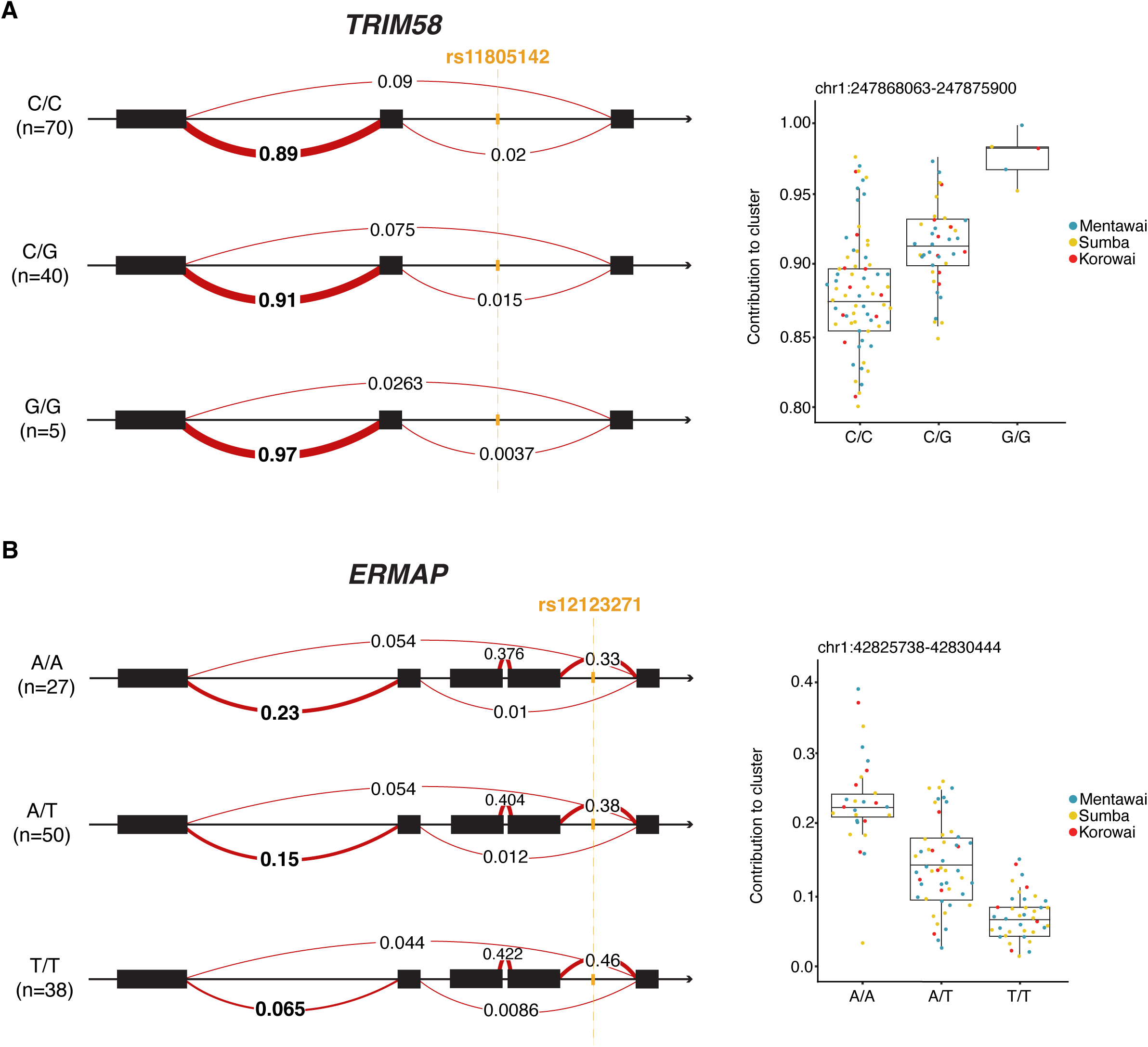
sQTL examples. **A)** Genotype plot of the *TRIM58* sQTL event. For this alternative splicing event, the junction with the most significant association to the SNP is highlighted with bold numbers across all three genotypes, and these numbers represent the proportion of reads spanning the junction. On the right, boxplots depict the distributions for each genotype. **B)** Genotype plot of the *ERMAP* sQTL event.

**Supplementary Figure 5.**
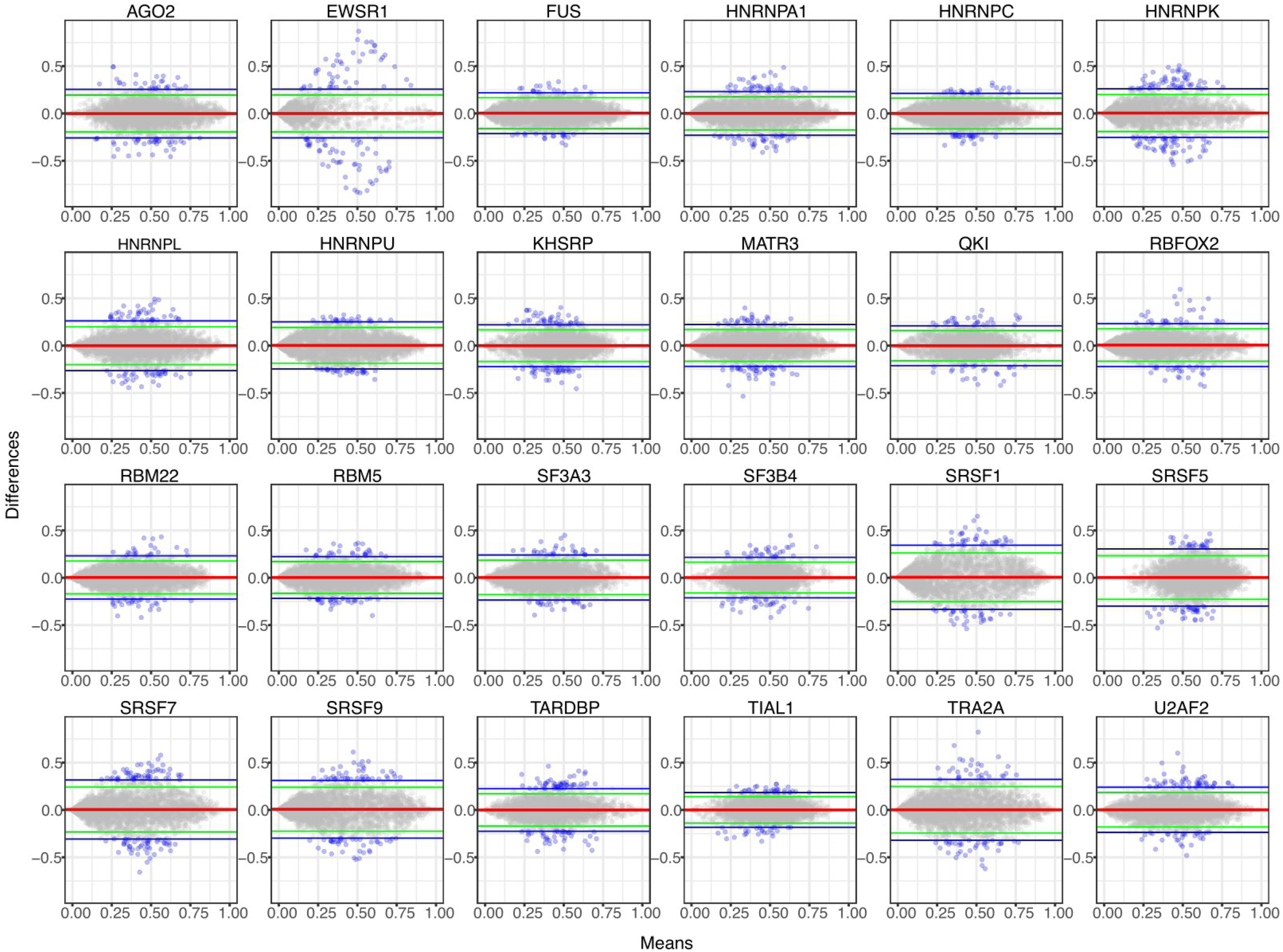
sQTL impact on RBP binding affinity. Mean-difference plots for 24 of the 33 splicing-RBPs. Each point represents an sQTL variant sequence (i.e., the sQTL with a 10bp flanking sequence) and reference sequence pair. The y-axis represents the difference between the DeepCLIP reference sequence score and the variant sequence score, while the x-axis depicts the average between these two scores. Horizontal green and blue lines represent the 95% and 99% CIs, respectively. Points that are outside of the 99% CI are colored blue.

**Supplementary Figure 6.**
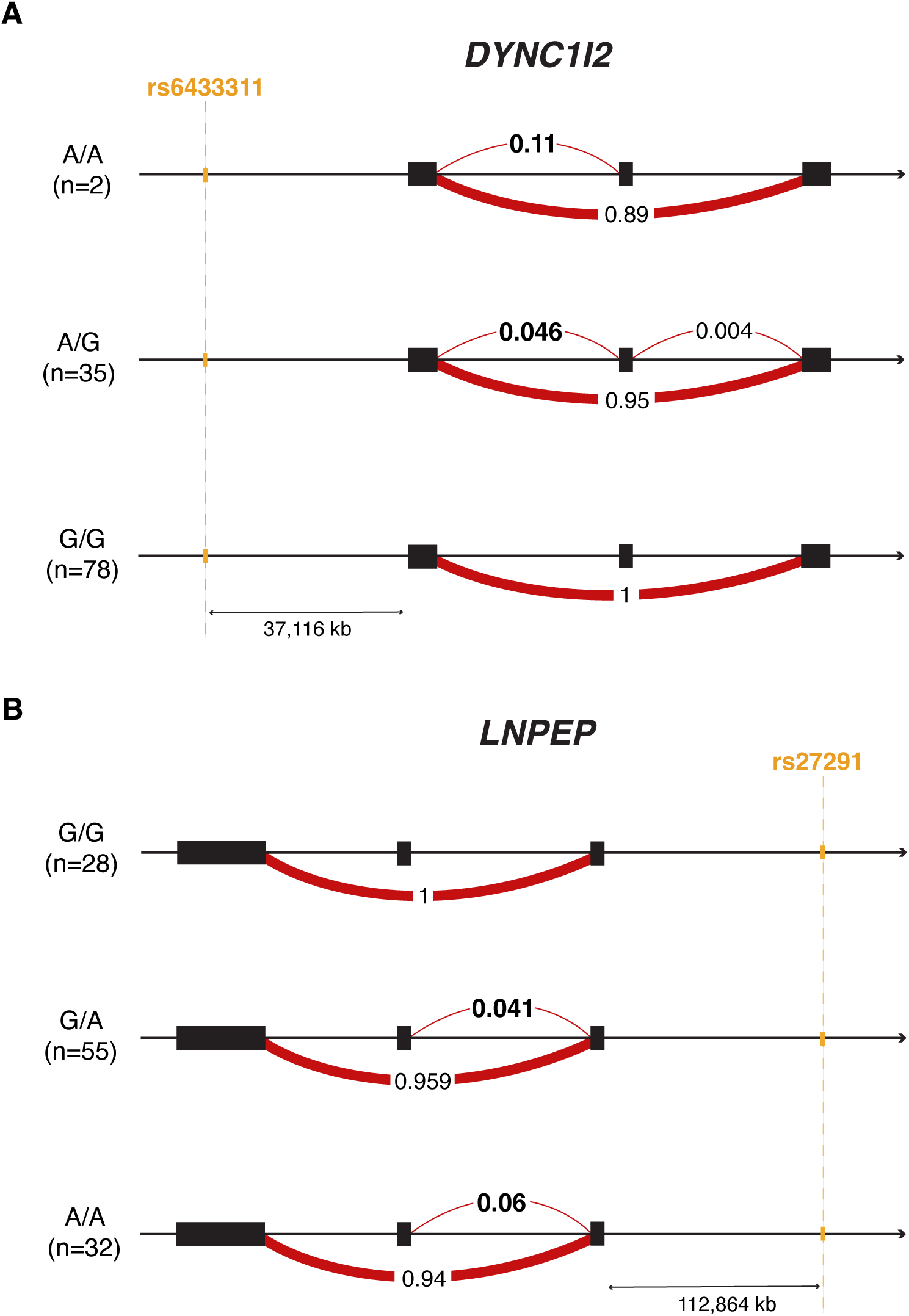
sQTL genotype plots for *DYNC1I2* and *LNPEP*. **A)** Genotype plot of the *DYNC1I2* sQTL event. For this alternative splicing event, the junction with the most significant association to the SNP is highlighted with bold numbers across all three genotypes, and these numbers represent the proportion of reads spanning the junction.

**Supplementary Figure 7.**
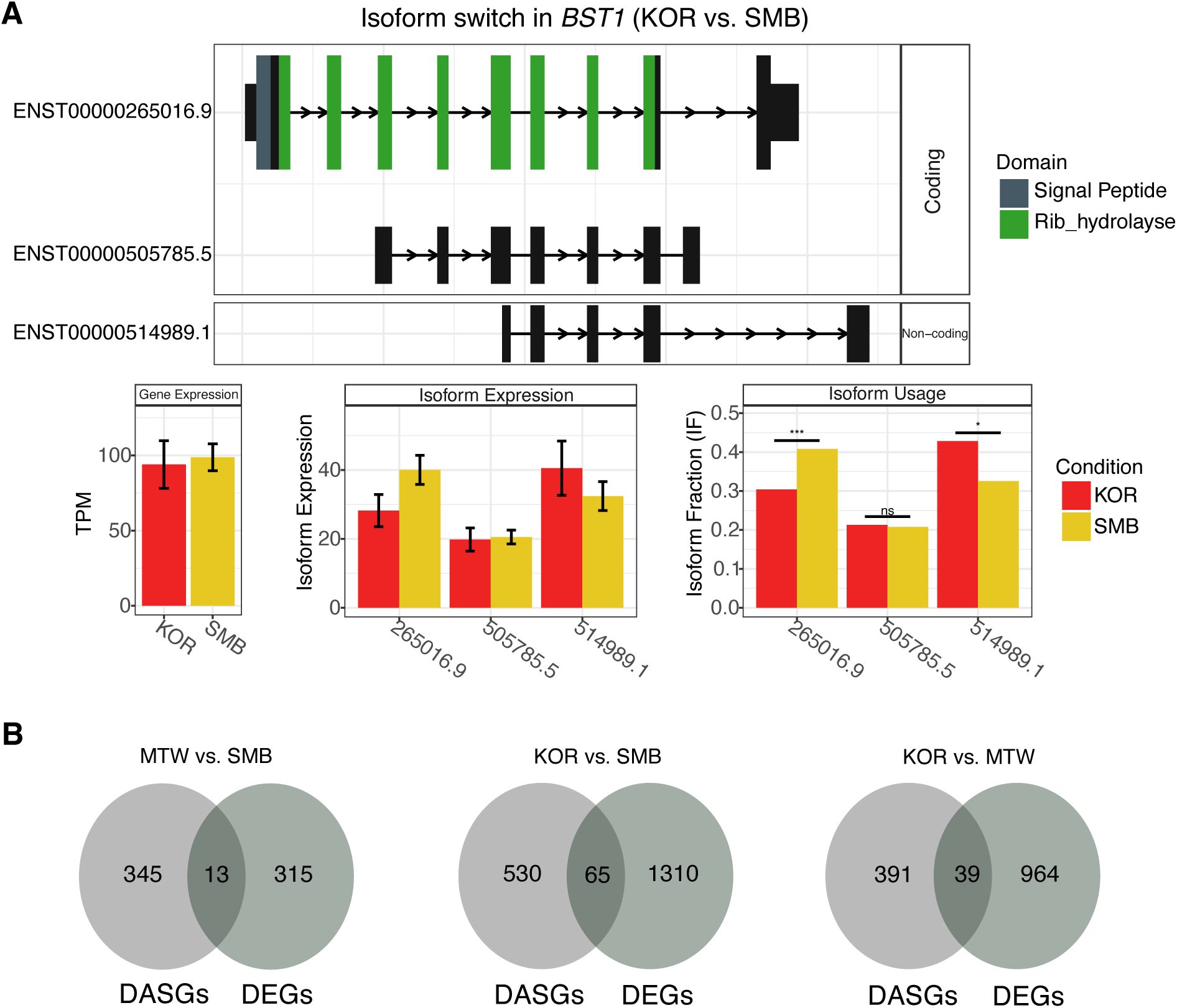
Comparing isoform expression and gene expression. **A)** Isoform switching analysis for *BST1* (between the Korowai population and the Sumba population). Exons with annotated domains are coloured, and those without are black boxes. Gene expression (TPM), as well as isoform expression and isoform usage are plotted for all detected isoforms. Error-bars indicate 95% confidence intervals. Level of significance for differences in isoform fraction (IF) is denoted by the asterisks. **B**) The overlap between differentially expressed genes (DEGs) and differentially alternatively spliced genes (DASGs).

## Supplementary Tables

**Table S1**

Gene Ontology and KEGG overrepresentation analysis for all AS genes.

**Table S2**

Gene Ontology and KEGG overrepresentation analysis for population-specific AS genes.

**Table S3**

Gene Ontology and KEGG overrepresentation analysis for DAS genes.

**Table S4**

Permutation-significant sQTLs. Available on Figshare: https://melbourne.figshare.com/projects/Profiling_genetically_driven_alternative_splicing_across_the_Indonesian_Archipelago/204111

**Table S5**

Nominal sQTL statistics. Available on Figshare: https://melbourne.figshare.com/projects/Profiling_genetically_driven_alternative_splicing_across_the_Indonesian_Archipelago/204111

**Table S6**

Gene Ontology and KEGG overrepresentation analysis for sGenes.

**Table S7**

sQTL-LAI correlation results for Papuan-like genetic ancestry.

**Table S8**

ClinVar annotation of lead sQTL SNPs.

**Table S9**

Summary of isoform switch analyses.

**Table S10**

Exon coordinates (GRCh38 Ensembl release 110) for all isoforms involved in switching events.

## Supplementary Files

**File S1**

sQTL-GWAS colocalization results.

**File S2**

CPC2 isoform coding potential prediction.

**File S3**

IUPred2A prediction of Intrinsically Disordered Regions (IDRs).

**File S4**

SignalP prediction of signal peptides.

**File S5**

PFAM domain prediction.

